# The Need for Transfer Learning in CRISPR-Cas Off-Target Scoring

**DOI:** 10.1101/2021.08.28.457846

**Authors:** Pavan K. Kota, Yidan Pan, Hoang-Anh Vu, Mingming Cao, Richard G. Baraniuk, Gang Bao

**Author notes:** These authors contributed equally to the paper (Pavan K. Kota, Yidan Pan).

## Abstract

**Motivation:** The scalable design of safe guide RNA sequences for CRISPR gene editing depends on the computational “scoring” of DNA locations that may be edited. As there is no widely accepted benchmark dataset to compare scoring models, we present a curated “TrueOT” dataset that contains thoroughly validated datapoints to best reflect the properties of *in vivo* editing. Many existing models are trained on data from high throughput assays. We hypothesize that such models may suboptimally transfer to the low throughput data in TrueOT due to fundamental biological differences between proxy assays and *in vivo* behavior. We developed new Siamese convolutional neural networks, trained them on a proxy dataset, and compared their performance against existing models on TrueOT.

**Results:** Our simplest model with a single convolutional and pooling layer surprisingly exhibits state-of-the-art performance on TrueOT. Adding subsequent layers improved performance on a proxy dataset while compromising performance on TrueOT. We demonstrate improved generalization on TrueOT with a Siamese model of higher complexity when we apply transfer learning techniques. These results suggest an urgent need for the CRISPR community to agree upon a benchmark dataset such as TrueOT and highlight that various sources of CRISPR data cannot be assumed to be equivalent.

**Availability and Implementation:** Our code base and datasets are available on GitHub at github.com/baolab-rice/CRISPR_OT_scoring.

## 1 Introduction

CRISPR-Cas9 systems are engineered for site- and sequence-specific genome editing [1, 2]. The *S. pyogenes* Cas9 (SpCas9) system is the most common variant and typically requires a 20-base guide RNA (gRNA) that targets a DNA sequence upstream of a Protospacer Adjacent Motif (PAM) sequence of “NGG” [3, 4]. The relatively high efficiency of SpCas9 systems and ease of construction in performing gene editing have led to a revolution in life sciences and medicine. However, unintended editing at “off-target” (OT) DNA sites is a major concern for gene editing applications [3]. The hybridization of the gRNA to target DNA tolerates imperfect sequence homology, and this can cause OT activity at DNA sites adjacent to a PAM. [5–7]. The resulting double strand breaks (DSBs) can induce undesired mutations that vary gene expression levels or even disrupt genes, and multiple DSBs can result in chromosomal rearrangement or severe DNA damage [8,9]. Therefore, rational designs of gRNAs with minimal OT activity are critical for both scientific studies and safe therapeutic applications of CRISPR-Cas9 systems.

As experimentally screening target DNA sites for potential OT activity is tedious and expensive, computational techniques are critical to scalably evaluate gRNA designs [10]. This evaluation has three phases for a given gRNA: screening for potential OTs across the whole genome, scoring the list of targets, and aggregating the scores into an overall gRNA quality metric [11]. In this work, we focus on scoring. Scoring models initially used hypotheses on relevant sequence features that affect editing activity [12–15]. Increased availability of CRISPR gene-editing data has enabled machine learning (ML) to directly learn relevant features from data [16]. Given a gRNA-target sequence pair as input, models predict a label that is either binary for classification (predicting whether a gRNA triggers CRISPR-Cas9 based editing at a target site) [17–19] or continuous for regression (predicting editing efficiency) [11, 20]. With few exceptions [19, 20], most scoring algorithms focus only on mismatches between the gRNA and target sequences without accounting for bulges. However, CRISPR-Cas9 systems have been shown to generate DSBs at sites with bulges in both *in vitro* and *in vivo* settings among multiple cell types [6, 21, 22].

Training a ML scoring model capable of assessing gRNA-target pairs with bulges is essential, but the available datasets are fundamentally limited. Ideally, datasets would label pairs based on the gRNA’s tendency to trigger an editing event at the target site *in vivo*, but such validation is painstaking. Whole-genome sequencing is limited by the sequencing depth and is generally unable to detect OT sites with less than 5% editing efficiency [23–25]. Therefore, many ML approaches rely on datasets generated by genome-wide high throughput methods based on proxies such as the insertion rate of a double-stranded DNA tag (GUIDE-seq [26]) or the cleavage rate *in vitro* (CIRCLE-seq [27], CHANGE-seq [22]) and *in vivo* (DISCOVER-seq [28], HTGTS [29]). Unfortunately, recent studies have shown that proxy assays have low concordance among each other and with validated *in vivo* editing [22, 30]. Perhaps due to this variable performance between assays, there is no gold standard benchmark dataset on which to compare scoring models despite the pervasive use of such benchmarks in other applications of ML. A carefully selected benchmark is urgently needed to guide the research of ML applications in CRISPR based editing.

Therefore, our first contribution is the curation of a novel benchmark “TrueOT” dataset. TrueOT contains 1903 binary-labeled datapoints that were thoroughly validated by mutation rates *in vivo*. We argue that *in vivo* editing prediction on gRNAs that are not seen during model training is the best performance metric for any scoring model. The use of proxy datasets to train models implicitly assumes that interactions between gRNAs and DNA are independent of the biological setting. Our second contribution is the unraveling of this assumption through the evaluation of a suite of Siamese convolutional neural networks. As we will discuss, this architecture is particularly helpful for testing the assumption of dataset equivalence through the lens of *transfer learning*. A core principle of transfer learning with neural networks is that initial convolutional layers capture features that generalize well between datasets while deeper layers must be retuned accordingly on a portion of the target dataset [31]. We trained the Siamese networks on a “Proxy Dataset” and found that our “S1C” model with a single large convolutional and pooling layer achieved state-of-the-art generalization to TrueOT among bulge-capable models. We found that more complex Siamese models improved performance on the Proxy Dataset while compromising generalization to TrueOT. We believe a transfer learning framework explains these results.

Our core thesis is that if TrueOT is an acceptable benchmark dataset, then future efforts in scoring model development should consider applying transfer learning principles of ML to account for the underlying gap between proxy assays and *in vivo* editing behavior. Researchers have similarly used transfer learning in model development to account for differences in gRNA activity in eukaryotes and prokaryotes [32]. TrueOT currently contains too few datapoints to train deep networks on directly. Still, we perform a preliminary demonstration of transfer learning through a novel dimensionality reduction on the output of our S1C to enable the tuning of a much smaller network. Our TrueOT benchmark and Siamese models serve as potent starting points for continued research into this problem. As more data is added to TrueOT, the efficacy of transfer learning will naturally improve along with the confidence in the safety of designed gRNAs.

## 2 Methods

### 2.1 Dataset Curation

#### 2.1.1 TrueOT

Currently, no assay enables direct genome-wide measurement of CRISPR-induced DNA sequence alteration with high sensitivity and throughput, significantly limiting the size of TrueOT. We defined three criteria for the inclusion of experimental data in TrueOT: (1) the experiments were performed in living cells, retaining the information that is missing from *in vitro* settings; (2) the OT editing efficiencies were evaluated by directly measuring the target sequence mutation rate, the common standard in clinical settings; (3) the OTs have a chromosomal position provided by original studies or a unique chromosomal position that can be retrieved in the reference genome hg38 using COSMID [13]. This filtering results in 1903 unique datapoints with 35 unique gRNAs from 11 different studies. Ten studies’ datapoints were experimentally validated through next-generation sequencing of PCR amplicons [9, 22, 33–40]. We positively labeled gRNA-target pairs with an editing rate greater than 0.1%, a commonly accepted threshold for deeming OTs [35, 41]. Although some scoring models use continuously valued editing efficiency for regression, we suggest that in evaluating potential OTs, any degree of editing may be dangerous and should be flagged accordingly. One study experimentally validated datapoints by T7 Endonuclease I for which we used the original study’s labels [5]. In determining which datapoints have bulges, we used the original studies’ alignment information to avoid adding a source of variability (Fig. S1). Among the 281 positive OTs in TrueOT, 10 have bulges, highlighting the need for bulge-aware scoring models. For further details on the included studies, see Table S1.

In our uploaded dataset (Table S2), we decided to keep all 1903 available datapoints that were performed in unique studies or unique experimental conditions. Some datapoints in TrueOT have the same gRNA-target pair but may exhibit different labels due to commonly observed cell-type dependencies in gene editing [5, 35, 42]. In all, there are 1786 unique gRNA-target pairs. Of these, 105 pairs have multiple entries with twenty pairs having conflicting labels. In model evaluation, we filtered for the 1806 unique triplets of {gRNA, target, label} to characterize the aggregate ability of models to make predictions on sequence information alone and reduce any ambiguous gRNA-target pairs to one instance of each a positive and negative label. Table S3 includes this duplicate removal in its description of TrueOT and the other datasets discussed below.

#### 2.1.2 Proxy Dataset

For training our models, we first combined a dataset from a recent review [43] and the training set from CRISTA [20]. The former includes datapoints from several publications but lacks datapoints with bulges, motivating the inclusion of the latter. We then excluded datapoints with gRNAs in TrueOT from this combined dataset, ensuring that our models’ performance on TrueOT reflects their ability to make *in vivo* editing predictions on unseen gRNAs. After this filtering, our Proxy Dataset has 3481 remaining datapoints from proxy assays, predominantly GUIDE-SEQ and HTGTS (Table S3).

### 2.2 Model Training and Validation

Figure 1a overviews how each dataset was used in the development and evaluation of our models. We trained and validated our own models exclusively on the Proxy Dataset. We built our models with Python 3.8.0 using TensorFlow 2.4.1. Throughout this work, splitting data is an approximate process due to the need to keep gRNAs unique in each subset of data. We trained three-fold cross validation (CV) ensembles on approximately 80% of the Proxy Dataset (“Proxy TrainCV”). Within each fold, we applied an inverse class weight to account for class imbalance and used early stopping based on CV performance. The ensembling process helps each member model learn to generalize to a different subset of gRNAs in Proxy TrainCV. The ensembles’ decisions were based on majority vote, and their ROC-AUC on the remaining 20% of data (“Proxy Validation”) guided hyperparameter selection. The number and width of convolutional filters as well as the number of neurons in dense layers were the primary hyperparameters that were optimized. The decision to use Proxy Validation to tune our models reflects the common practice of using generalization on proxy assays as a ranking mechanism of scoring algorithms.

**Figure 1:**
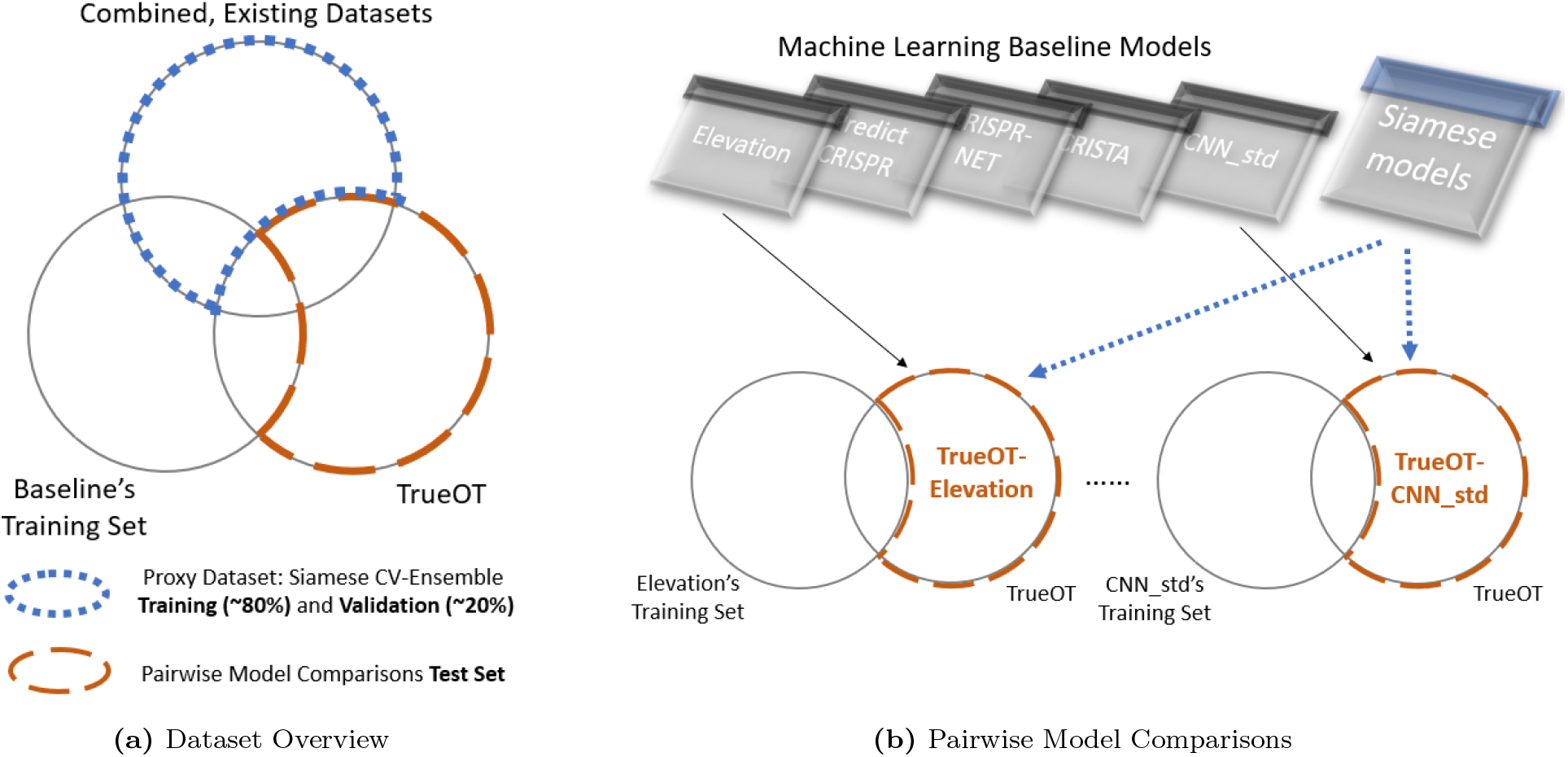
Schematics for how we used each dataset. (a) This overview applies to all Siamese models reported in this work except for the S1C_Hybrid_TL (Sections 2.4.4 and 3.5). To generate our internal Proxy Dataset for developing the Siamese models, we combined two existing datasets and filtered them to exclude datapoints with gRNAs in TrueOT. We split the Proxy Dataset into a TrainCV portion for training ensembles and a Validation portion for hyperparameter selection. TrueOT served as an external test set for Siamese model comparison and pairwise comparisons against existing models. (b) Each machine learning baseline model was evaluated on the subset of TrueOT that excluded datapoints with gRNAs found in the baselines’ training sets. The subset of TrueOT for comparison is thus a function of the baseline model.

### 2.3 Pairwise Model Comparisons on TrueOT

We tested our models on TrueOT and compared their performance against that of existing algorithms. In addition to several rule-based models [6, 10, 13–15, 44, 45], we chose the following recent ML models for their high performance as evaluated by a recent review [43]: CRISTA [20], Elevation [11], predictCRISPR [18], CNN std [17]. We also include CRISPR-NET [19], a recently developed bulge-aware scoring model. To evaluate the baseline models as published on TrueOT, we do not have the luxury of retroactively removing datapoints in the original training sets that contain gRNAs in TrueOT. Retraining existing models on new datasets risks introduction of bias or errors due to our own implementation, so instead, we evaluated the performance of the published models on subsets of TrueOT involving gRNAs that were not used in the training of the original model. We also filtered datapoints for which the model cannot produce an output such as a bulge-containing datapoint with a mismatch-only algorithm. Figure 1b illustrates this process, and Table S3 provides a quantitative breakdown of the subsets of TrueOT for all baseline models.

For each baseline model, we evaluated the area under the curve of the receiver operating characteristic and precision-recall curves (ROC-AUC, PR-AUC). Notably, Elevation and CRISTA are regression models whereas we compared classification performance. Sweeping thresholds in the evaluation of ROC-AUC and PR-AUC can standardize such comparisons. We evaluated our own model architectures using five different initial random seeds and performed a one sample *z*-test relative to the AUCs of the baselines.

### 2.4 Siamese Model Design

As our core architecture, we selected a Siamese network which is commonly used for sequence comparisons in natural language processing with some recent adaptations to biological settings [46, 47]. A Siamese network passes elements of paired data through an identical sequence processing network and evaluates their similarity. Here, we assessed the Euclidean distance between the gRNA and target sequences’ network output. By optimizing a contrastive loss function, Siamese networks learn to position similar sequences (i.e., gRNA-target pair that results in editing) close together while pushing dissimilar pairs further apart.

A Siamese network is particularly helpful for investigating transfer learning via the influence of various network depths because its output dimension is arbitrary; layers can easily be added or removed without having to condense the final output to a scalar value (regression) or the number of classes (classification). Such dimensionality reduction in other neural networks is often accomplished by dense layers which are parameter intensive and unlikely to transfer well between datasets.

#### 2.4.1 Data Encoding

Existing bulge-aware scoring algorithms require alignment for data encoding [19, 20]. Because alignment algorithms can vary in output given the same pair of sequences, scoring algorithms may be susceptible to these disagreements. We simplified the encoding strategy to ignore exact alignment and attempted to correct for small frameshifts caused by bulges through a max pooling operation. The gRNA and target nucleotides {A, C, G, T/U} are one-hot encoded, and by simply left-padding the sequences with zeros to a fixed length of 26, our encoding is alignment-independent (Fig. 2a). We chose 26 as up to three insertions have been allowed in past alignment techniques for CRISPR scoring [20].

**Figure 2:**
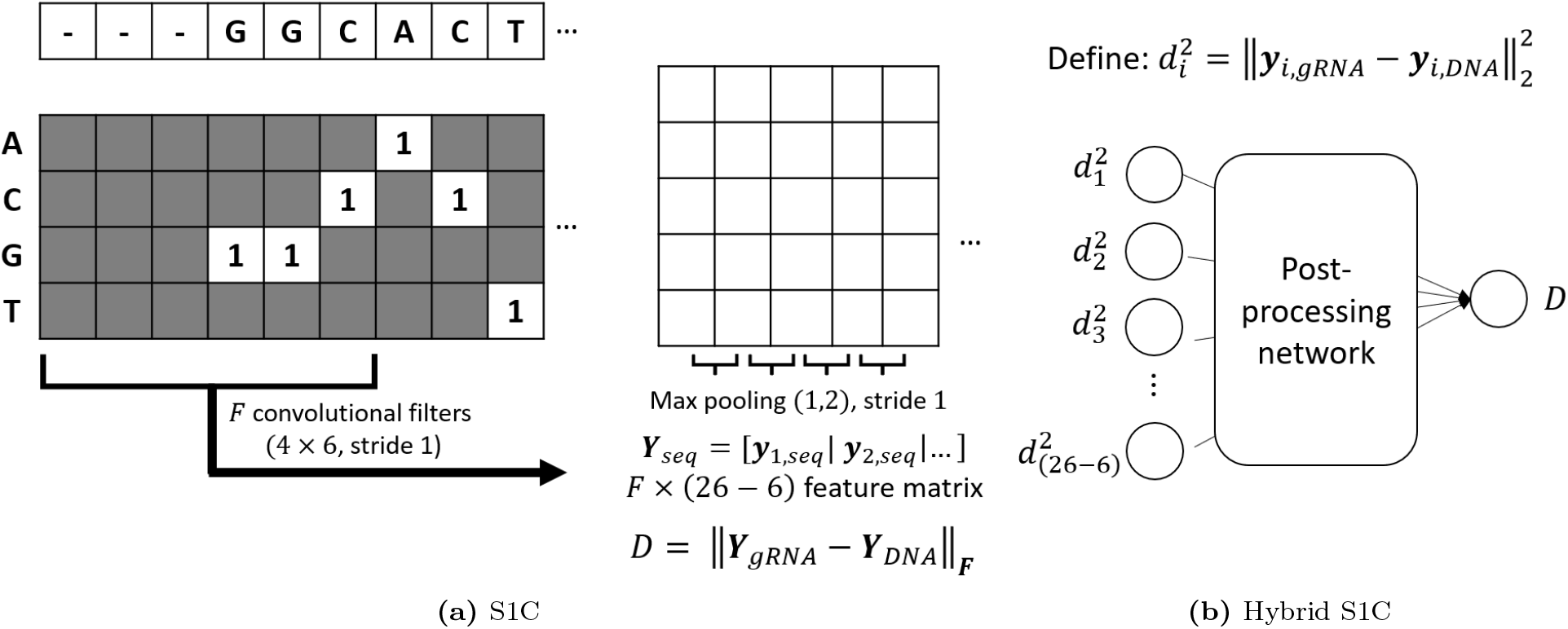
The Siamese models compute a distance *D* between a guide RNA (gRNA) and target DNA sequence. (a) Rather than aligning the gRNA and target sequences, we simply left-pad them to a fixed length of 26. The S1C applies a single convolutional and pooling layer to each one-hot encoded sequence. The filter vector at position *i* for a sequence *seq* is denoted *y*_*i,seq*_. The distance is the standard Euclidean distance between the feature matrices of the gRNA and DNA sequences. (b) The Hybrid S1C computes the squared distance between two input sequences at each position from the output of an S1C. Note that the S1C’s distance can be represented by 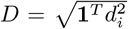. By passing the position-wise distances to a post-processing network with a nonnegative activation function (e.g., ReLU), the Hybrid S1C can learn a nonlinear “distance” guided by a contrastive loss function. The post-processing network is entirely arbitrary in principle, although we use a single dense layer of 128 neurons in this work.

#### 2.4.2 Siamese Networks

We started with a single convolutional and pooling layer with many filters because of such layers’ established ability to capture features that generalize across datasets [31]. This Siamese 1-Convolution model (S1C) anomalously allows filters to be added indefinitely without harming generalization (in theory) since Siamese models’ output dimension can be arbitrarily large and a distance is immediately computed after pooling. As a result, we used 2^13^ = 8192 filters of six nucleotide width in our S1C based on our hyperparameter search. For bulge awareness, we added a 1 *×* 2 max pooling operator with stride 1, allowing our network to partially tolerate *±*1 position frameshifts of sequences. We also hypothesized that our relatively wide six-nucleotide convolutions would learn some frameshift tolerance themselves.

Despite these features, there are several reasons why additional model complexity is warranted. First, the S1C only measures local dependencies within six nucleotide windows. By computing a distance on the output of this layer, neither positions nor combinations of features are considered although both have established relevance [6, 13]. There are arbitrarily many ways to increase model complexity, but we present just two simple cases representative of the evidence of a need for transfer learning. In one implementation, we added a second convolutional and pooling layer (S2C) with eight filters, and in another, we added a dense layer of 32 neurons to the S2C (S2C_Dense). For these two extensions, we constrained the number of filters (32) in the first convolutional layer to prevent an explosion of parameters. To directly compare S2C and S2C_dense against a single convolutional layer, we defined the S1C_mini model with just these 32 filters.

#### 2.4.3 Untrained S1C

Random filter weights in convolutions have been shown to create distance-preserving (“isometric”) embeddings of inputs [48, 49], making them a natural choice for a Siamese network that makes decisions based on the output positions of input sequences. Before convolution, the distance between a one-hot encoded gRNA and OT is scaled by the number of mismatches, meaning an untrained S1C’s filters will approximately count mismatches in each six nucleotide window. Training filter weights distorts the network’s output positions to better separate classes, especially with datapoints at the boundary [49]. In the context of CRISPR scoring, boundary points are gRNA-target pairs with strong sequence homology that do not result in editing and vice versa. We compared the performance of an untrained S1C model (“S1C_ut”) against that of the trained Siamese models and existing baselines to provide insight into the influence of training on the resolution of boundary points.

#### 2.4.4 Hybrid S1C Models for Transfer Learning

A common technique in transfer learning is to freeze the weights of high level convolutional layers from a model pre-trained on a larger dataset, add subsequent layers to the network, and train only the subsequent layers’ weights on a portion of the target dataset. In our demonstration of transfer learning, we used the S1C pre-trained on Proxy TrainCV as the initial model, but we must reasonably control the complexity of subsequent layers since TrueOT is a very small target dataset. The raw output dimension of the S1C is very large, making the immediate application of a subsequent layer parameter intensive. However, the S1C is position-invariant with an output equivalent to the square root of the sum of the squared position differences in each convolutional window. This intuition lends a natural way to dramatically reduce the output dimension of the S1C for transfer learning on a small dataset: we compute the squared position differences and pass this small vector to a dense network (Fig. 2b). With a ReLU activation on the final output, this “Hybrid S1C” learns a nonlinear “distance” metric that accounts for combinations of positions of sequence discrepancies between the gRNA and target. We remove the square root function on the output since it is a monotonic function and ROC and PR characterizations are based on thresholds.

For the “S1C_Hybrid_TL,” we split TrueOT approximately into 75% (TrueOT TrainCV) for training and CV and 25% (TrueOT Test) for testing, enforcing non-overlapping gRNAs between each split. We performed a similar three-fold CV training and ensembling process described in Section 2.2. For each fold of the S1C_Hybrid_TL, we loaded the weights of the corresponding fold from the original S1C and froze them, trained only the additional dense layers of the post processing network, and used the CV portion to guide early stopping. We used CV performance for hyperparameter selection, ultimately choosing a single hidden layer of 128 neurons. This process maintained the S1C_Hybrid_TL as a three-member ensemble.

For the “S1C_Hybrid_Proxy,” we trained the entire network including the convolutional layer on Proxy TrainCV using the same hyperparameters. Among multiple initializations, we selected that with the best Proxy Validation performance. We generated ten different splits of TrueOT into TrueOT TrainCV and TrueOT Test and evaluated the change in ROC-AUC performance on TrueOT Test relative to the original S1C. Repeated splitting of TrueOT ensures that any apparent performance change is not just an anomaly of a particular split. Comparing these two versions of the Hybrid S1C indicates whether the performance change is due to the transfer learning process specifically or merely due to the change in network architecture.

## 3 Results

### 3.1 Model Design by Proxy Validation Compromises TrueOT Performance

Table 1 illustrates a general pattern from our research: improving generalization performance on the Proxy Validation set can compromise the performance on TrueOT. In Table 1, Split 1 refers to the random split of the Proxy Dataset used to train and select hyperparameters of our Siamese models. Recall that S2C and S2C_dense are built off of the S1C_mini. These simple extensions generally improve performance on the Proxy Validation set while exhibiting worse performance on TrueOT than S1C_mini. For the same model architectures, we repeated the training and evaluation process for four other splits of the Proxy Dataset to ensure the pattern was not anomalous of Split 1. Indeed, the relative performances generally hold and the Spearman rank correlation (*r*_*s*_) between Proxy Validation and TrueOT performance is consistently negative. We speculate that *r*_*s*_ has the highest magnitude in Split 1 as this data split was used for designing the models towards improved Proxy Validation performance.

**Table 1:** Performance (ROC-AUC) of Siamese models on various splits of the Proxy Dataset. Split 1 was used for hyperparameter selection for all models, and all model architectures were retrained on four other random splits of the Proxy Dataset. Namely, Proxy TrainCV and Proxy Validation are a function of the split. Each Siamese model was trained with five different initial random seeds, and the results of the model whose seed achieved median Proxy Validation performance are shown here.

| | | S1C_ut | S1C | S1C_mini | S2C | S2C_dense | $r_s$ |
| --- | --- | --- | --- | --- | --- | --- | --- |
| Split 1 | Proxy Val. | 0.647 | 0.645 | 0.625 | 0.694 | 0.731 | -0.9 |
|  | TrueOT | 0.794 | 0.827 | 0.794 | 0.761 | 0.747 |  |
| Split 2 | Proxy Val. | 0.763 | 0.749 | 0.822 | 0.772 | 0.769 | -0.3 |
|  | TrueOT | 0.794 | 0.814 | 0.811 | 0.775 | 0.744 |  |
| Split 3 | Proxy Val. | 0.682 | 0.755 | 0.736 | 0.773 | 0.808 | -0.5 |
|  | TrueOT | 0.794 | 0.812 | 0.812 | 0.730 | 0.747 |  |
| Split 4 | Proxy Val. | 0.711 | 0.739 | 0.720 | 0.715 | 0.776 | -0.3 |
|  | TrueOT | 0.794 | 0.806 | 0.786 | 0.755 | 0.713 |  |
| Split 5 | Proxy Val. | 0.702 | 0.751 | 0.730 | 0.765 | 0.759 | -0.5 |
|  | TrueOT | 0.794 | 0.822 | 0.794 | 0.769 | 0.739 |  |

The untrained S1C_ut performed notably better on TrueOT than on various splits of the Proxy Validation set, meaning that classes of sequence pairs in TrueOT are more closely related to an absolute count of mismatches along with some frameshift tolerance. Although we have no influence over the distribution of data in TrueOT, this contrast illustrates that our Proxy Validation set has many more boundary points from which our Siamese models may hope to learn. Concerningly, the S1C_ut consistently outperformed the S2C and S2C_dense on TrueOT despite underperforming them on Proxy Validation. Any boundary rules that the more complex models learned from Proxy TrainCV appear to have been misleading for TrueOT.

Ultimately, these results indicate that the Proxy Validation set is suboptimal for guiding model design if the goal is to generalize to TrueOT. We speculate that this is caused by an underlying discrepancy in the biology of proxy assays versus *in vivo* editing. If the two datasets were from the same underlying distributions, this inverse effect would be very unlikely to occur. Moreover, the single-convolutional Siamese models performed the best and comparably on TrueOT. This result is consistent with the established robustness of high level convolutional layers in transfer learning applications.

### 3.2 S1C Achieves State-of-the-Art Performance on TrueOT

We compared the S1C against baselines given the results in Section 3.1 and included S1C_ut as a useful reference. In pairwise comparisons on appropriate subsets of TrueOT, our S1C significantly outperformed most of the tested baseline algorithms (Table 2) with a few exceptions. Elevation and predictCRISPR outperformed the S1C, and CRISPRoff performed nearly identically to the S1C. We speculate that this similarity could be due to CRISPRoff’s primary reliance on the nearest-neighbor model for nucleic acid interactions which sum the thermodynamic effects of small windows of nucleotides across the duplex [45], a structurally similar approach to the S1C (Fig. 2). However, these models cannot account for bulges, meaning that the comparisons are restricted to gRNA-target pairs of equal length.

**Table 2:** Pairwise Comparisons on TrueOT. We conduct one-sided *z*-tests between the Baselines and the S1C’s. Baselines’ AUCs are bolded if they are greater than both the S1C’s and S1C_ut’s AUCs with *p <* 0.001. The S1Cs’ AUCs are bolded if they are greater than the Baselines’ AUC with *p <* 0.001.

|  |  | ROC-AUC |  |  | PR-AUC |  |  |
| --- | --- | --- | --- | --- | --- | --- | --- |
| Baseline Name | $n$ | Baseline | S1C | S1C_ut | Baseline | S1C | S1C_ut |
| Cropit | 1579 | 0.676 | **0.818** | **0.782** | 0.330 | **0.519** | **0.472** |
| Hsu | 1579 | 0.667 | **0.818** | **0.782** | 0.356 | **0.519** | **0.472** |
| CCTop | 1579 | 0.652 | **0.818** | **0.782** | 0.320 | **0.519** | **0.472** |
| MIT | 1579 | 0.758 | **0.818** | **0.782** | 0.421 | **0.519** | **0.472** |
| CFD | 1579 | 0.805 | 0.818 | 0.782 | 0.465 | **0.519** | **0.472** |
| COSMID | 1784 | 0.659 | **0.824** | **0.791** | 0.327 | **0.510** | **0.464** |
| CRISPRoff | 1481 | 0.817 | 0.818 | 0.781 | 0.546 | 0.553 | 0.505 |
| CNN_std | 1044 | 0.707 | **0.802** | **0.755** | 0.375 | **0.461** | **0.401** |
| Elevation | 1116 | 0.800 | 0.784 | 0.744 | **0.443** | 0.405 | 0.377 |
| predictCRISPR | 857 | **0.810** | 0.757 | 0.713 | **0.396** | 0.365 | 0.328 |
| CRISTA | 1252 | 0.731 | **0.781** | **0.743** | 0.354 | **0.384** | 0.355 |
| CRISPR-NET | 1078 | **0.779** | 0.758 | 0.715 | 0.198 | **0.328** | **0.293** |

Among all baselines, COSMID, CRISTA, and CRISPR-NET are designed to account for bulges. In aggregate, the S1C appears roughly on par with CRISPR-NET, slightly underperforming in ROC-AUC and outperforming in PR-AUC (Table 2). We further partitioned TrueOT into datapoints with and without bulges to deepen our comparison against these models (Table 3). The S1C significantly outperformed the baselines in almost all categories. The only exception appears to clarify the comparable overall performance of the S1C and CRISPR-NET in Table 2: CRISPR-NET had an edge on bulge-excluded datapoints while the S1C had better performance on bulge-containing datapoints.

**Table 3:** Pairwise Comparisons on TrueOT for bulge-capable models. The original *n* datapoints available for pairwise comparisons were split into bulge-containing gRNA-target pairs and all other pairs. A pair was considered to have a bulge if a ‘-’ appeared in either sequence in the original study’s alignment or if the two sequences were of different lengths. Bold font is applied as in Table 2.

| Baseline Name | TrueOT Subset | $n$ | ROC-AUC | | | PR-AUC | | |
| --- | --- | --- | --- | --- | --- | --- | --- | --- |
|  |  |  | Baseline | S1C | S1C_ut | Baseline | S1C | S1C_ut |
| COSMID | No Bulge | 1579 | 0.646 | **0.818** | **0.782** | 0.335 | **0.519** | **0.472** |
|  | Bulge | 205 | 0.733 | **0.832** | **0.813** | 0.279 | **0.382** | **0.345** |
| CRISTA | No Bulge | 1066 | 0.739 | **0.775** | 0.734 | 0.374 | **0.397** | 0.368 |
|  | Bulge | 186 | 0.584 | **0.822** | **0.806** | 0.089 | **0.383** | **0.346** |
| CRISPR-NET | No Bulge | 896 | **0.844** | 0.748 | 0.699 | **0.359** | 0.339 | 0.304 |
|  | Bulge | 182 | 0.643 | **0.818** | **0.802** | 0.073 | **0.383** | **0.346** |

The S1C’s approximate equivalence with the state-of-the-art should not be taken lightly. By including only a single convolutional layer, the S1C has no capacity to learn nonlinear combinations of features, unlike all of the ML-based methods noted here. Moreover, even the untrained S1C_ut outperformed most rule-based and some ML models on TrueOT, and it outperformed all baselines on datapoints with bulges. The S1C_ut essentially counts the number of mismatches with some frameshift tolerance. While more patterns are clearly necessary to distinguish boundary points in a dataset, it appears that the added functional capacity of existing models does not necessarily improve *in vivo* editing predictions.

### 3.3 Baseline Models’ Datasets Reflect their TrueOT Performance

Our pairwise model evaluations are intended to compare baselines against the S1C and do not directly reflect a rank ordering among baselines. However, the relative performance appears consistent with our transfer learning hypothesis; better performing baselines incorporate more *in vivo*-based data in their training process. PredictCRISPR used many of the low-throughput datapoints contained in TrueOT in its training set [18], hence its fewest datapoints *n* on which we could perform a pairwise comparison fairly. CFD performed the best among the rule-based algorithms on the full 1579 bulge-excluded datapoints and came very close to the performance of the S1C. We suggest that while CFD is often labeled as rule-based in the literature, it could be considered an ML approach driven by *in vivo* data as its weights are tuned based on flow cytometry data [6]. These direct *in vivo* measurements, while not based on sequence modification rate for inclusion in TrueOT, are arguably very close in motivation. Lastly, Elevation derived its model as a generalization of CFD, which may explain its similarly high performance. More details on the datasets used in each ML baseline are available in Table S4.

### 3.4 Bulge Performance of the S1C Appears Driven by Max Pooling

The S1C’s superior performance on the bulge-containing subsets of TrueOT (Table 3) could be due to chance given the small number of datapoints available (*n ≤* 205). Nonetheless, we investigated the decision-making process of the S1C for bulges to understand its high performance. We hypothesized that the use of many filters (2^13^) allowed the S1C to treat mismatches due to small frameshifts from bulges differently than random mismatches. For example, an insertion at position 4 in a window on the target sequence could appear as 3 mismatches with the gRNA (Fig. 3a), but perhaps the S1C learns to recognize such an occurrence as a frameshift instead.

**Figure 3:**
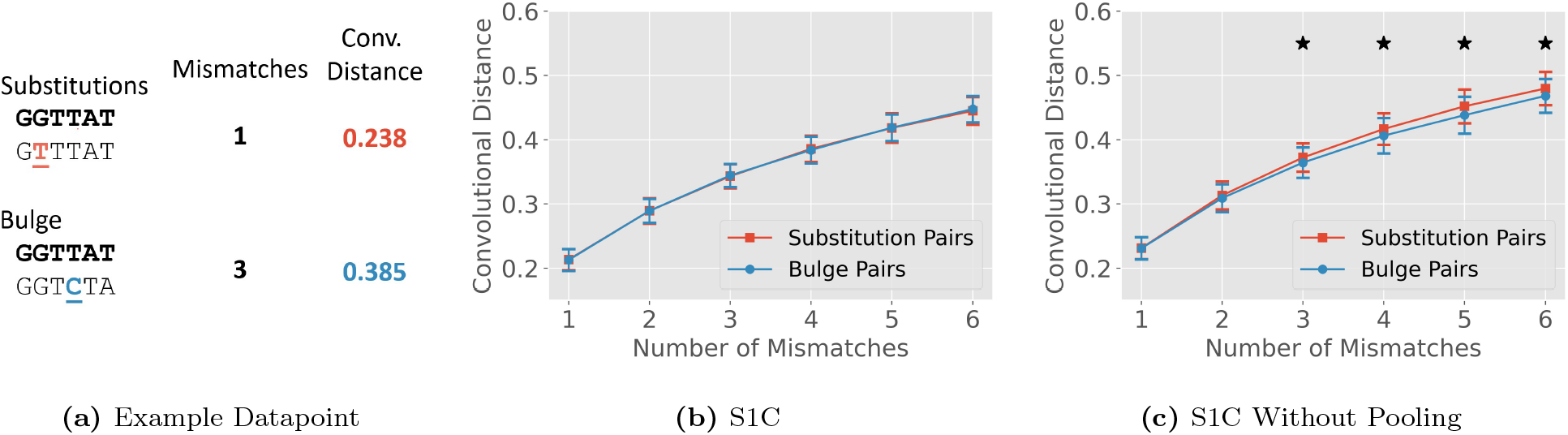
Characterization of the capacity of convolutions to learn the effect of bulges. Error bars *±*1 s.d. (a) Pairs of sequences were generated either by introducing substitutions or bulges at specific positions. For each pair in each method of mutation, the raw number of mismatches and convolutional distance was evaluated. (b) Comparison of the relationship between mismatches and network distance for the S1C. (c) Same comparison with an S1C without pooling. In (b) and (c), a black star indicates *p <* 0.001 in a two-sided Welch’s *t*-test.

We tested this hypothesis by generating a series of random sequences of six nucleotides and pairing them either with a mutated sequence with a random number of substitutions or with a sequence modified by a single-nucleotide insertion or deletion. To maintain a sequence length of six, the 3’ base was truncated for insertions; a random base was appended to the 3’ end for deletions. We evaluated the distances computed by the S1C’s filters between each pair of sequences and plotted them as a function of the naive mismatch count. While we expected the bulge-generated pairs to exhibit lower convolutional distances than the mismatch-generated pairs - an indication of recognizing greater similarity - we were surprised to find essentially the same distribution of distances for both sets of sequences (Fig. 3b). The distance computation between pairs appears unaware of bulges, indicating that the S1C is managing bulges predominantly through max pooling. Indeed, the untrained S1C_ut performs only slightly worse than the S1C on bulge datapoints (Table 3) and is *only* utilizing max pooling for bulge awareness. When we remove pooling and retrain the S1C with an otherwise identical network, the filters are forced to learn to recognize bulges on their own by considering mismatches due to bulges are considered as slightly “closer” than those caused by substitutions (Fig. 3c).

### 3.5 Transfer Learning with Hybrid S1C Improves TrueOT Generalization

In evaluating the effect on ROC-AUC for ten different splits of TrueOT Test (Fig. 4), we found that the S1C_Hybrid_TL has improved ensemble performance over the S1C (one-sided paired *t*-test, *p* = 0.04) and the S1C_Hybrid_Proxy (*p* = 0.02), the latter two of which were trained on Proxy TrainCV. The S1C_Hybrid_Proxy lends an insignificant improvement on TrueOT Test performance over the S1C (*p* = 0.34). These results corroborate our hypothesis that the source of training data influences generalization to *in vivo* behavior. Granted, these *p* values are smaller than the 0.001 significance level used in our tables. We speculate that the small improvement is mostly due to limited training data in splits of TrueOT TrainCV. Future research may see larger performance gains as more data is available for inclusion in TrueOT for tuning deeper layers.

**Figure 4:**
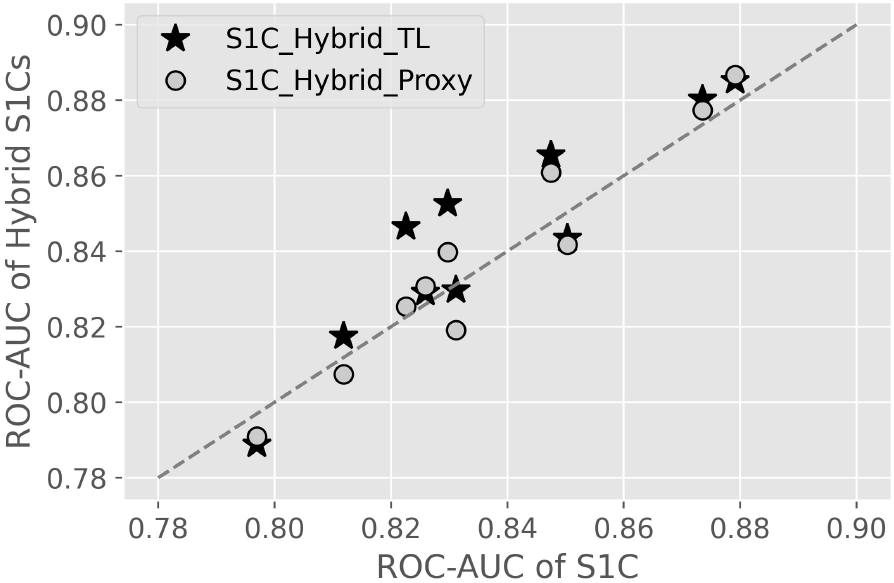
Performance comparison of the S1C_Hybrid_TL (stars) and S1C_Hybrid_Proxy (circles) against the original S1C (dashed line) on ten different splits of TrueOT Test. Vertically aligned stars and circles are evaluated on the same split of TrueOT Test. In one-sided paired *t*-tests, the S1C_Hybrid_Proxy elicits insignificant improvement over the S1C in TrueOT Test generalization (*p* = 0.34) whereas the S1C_Hybrid_TL lends performance greater than that of the S1C_Hybrid_Proxy (*p* = 0.02) and the original S1C (*p* = 0.04).

## 4 Discussion

The computational scoring of putative OTs is essential for economically designing safe gRNAs. Scoring models to date have reported test set performance on datasets sourced by assays that measure a proxy of *in vivo* editing. Each of these sources carries the risk of being misaligned with true *in vivo* editing behavior. Models should be evaluated based on their ability to predict the *in vivo* editing behavior of gRNA sequences that were unseen during model training [50]. We suggest that our curated TrueOT dataset offers a starting point for an accepted benchmark, and our initial exploration towards a new model supports our hypothesis of an underlying mismatch between various datasets.

Our simple S1C model with only one convolutional layer achieved state-of-the-art performance on TrueOT as reflected by pairwise comparisons against several rule-based and ML models. This result suggests the potency of a Siamese convolutional architecture for deriving generalizable features across datasets. Moreover, adding subsequent layers to S1C improved generalization performance on our Proxy Validation set while compromising performance on TrueOT. This inverse relationship strongly suggests that the data in the two datasets originate from different distributions; sequence features that govern editing *in vivo* appear to be sufficiently distinct from those of the assays represented in Proxy Validation.

We are concerned that an underlying discrepancy among datasets is a pervasive issue in the CRISPR scoring literature. Notably, even the untrained S1C_ut performed better than many baseline models on TrueOT and achieved state-of-the-art performance on datapoints with bulges. This result indicates that baselines may have learned irrelevant (if not misleading) features for TrueOT during training on data from proxy assays, an effect we observed during our own development of Siamese networks (Table 1). While researchers commonly use one or a few external datasets for model comparisons, there is little consideration of which dataset is more reflective of *in vivo* editing. Therefore, we believe there is an urgent need for a common benchmark dataset such as TrueOT to compare models. We acknowledge that this work does not consider the influences of epigenetic features that can lend variable behavior between cell types. As more *in vivo*-validated data becomes available, we envision TrueOT being split into nuanced subsets of data for particular applications.

If training data is sourced from proxy assays, transfer learning principles should be considered for improving performance on such benchmark datasets. Our S1C’s high performance and extreme simplicity offers a strong baseline against which to compare new models and a potent starting point for extended research as there is substantial room for improvement. A single convolutional layer aggregates local relationships without considering position or nonlinear combinations of features. Added complexity directly to the S1C as posed here will be difficult with its abnormally high (2^13^) number of filters (an idiosyncrasy of the S1C architecture). However, either the S1C_mini with far fewer (2^5^) filters or the dimensionality reduction offered by the Hybrid S1C framework could serve as more accessible starting points. In any case, our results suggest that benefiting from additional model complexity will be increasingly feasible as data is added to TrueOT and transfer learning approaches are considered.

We are optimistic that the expansion of TrueOT and applied transfer learning principles will help alleviate other issues in CRISPR datasets by better elucidating the causes of OT activity. For instance, it is widely assumed that the targets with low sequence homology to a gRNA will have zero editing efficiency such that in many assays, signals from such target sites are treated as noise and excluded from the final output [26,51]. These exclusions may include datapoints with bulges depending on the particular alignment method used to gauge sequence homology [20]. While the field gradually recognizes the importance of OTs with bulges, the noise filtering process based on homology carries the risk of mislabeling target sites with bulges. Further experimental validations on sites with low sequence homology will help distinguish genuine OTs from noise generated by somatic mutations and DNA repairs while also benefiting the basic research of gRNA-target hybridization mechanism.

In conclusion, there is an unmet need for a widely accepted gold standard dataset for benchmarking OT evaluation pipelines. While we propose the TrueOT dataset and its corresponding inclusion criteria, we urge the CRISPR community to more broadly recognize the need for such a dataset and modify TrueOT as it sees fit. We find substantial evidence that the datasets used in model development should not be considered equivalent from a machine learning perspective. Instead, they appear to have discrepancies such that the decision-making processes learned from one dataset may transfer suboptimally to another. As more experimentally validated *in vivo* editing data becomes available, dedicated transfer learning efforts can begin to properly leverage the quantity of high-throughput data.

## Acknowledgments

This work was supported by the National Institutes of Health [UG3HL151545, R01HL152314 to G.B.]; the Office of Naval Research Vannevar Bush Faculty Fellowship [N00014-18-1-2047 to R.G.B.]; and the National Library of Medicine Training Program [T15LM007093 to P.K.K.].

## Notes

https://github.com/baolab-rice/CRISPR_OT_scoring

### Competing Interest Statement

The authors have declared no competing interest.

### Summary of Updates

Fixed small mistake in original Proxy Validation dataset; Siamese models redesigned based on updated validation dataset, resulting in minor changes; All Figures and Tables revised accordingly; New Figure 1 to clarify delineations between datasets; New Table 1 replaces original Figure 2; New Table S3 adds detail on the composition of the datasets.

