## Supplemental Figures for "The Need for Transfer Learning in CRISPR-Cas Off-Target Scoring"

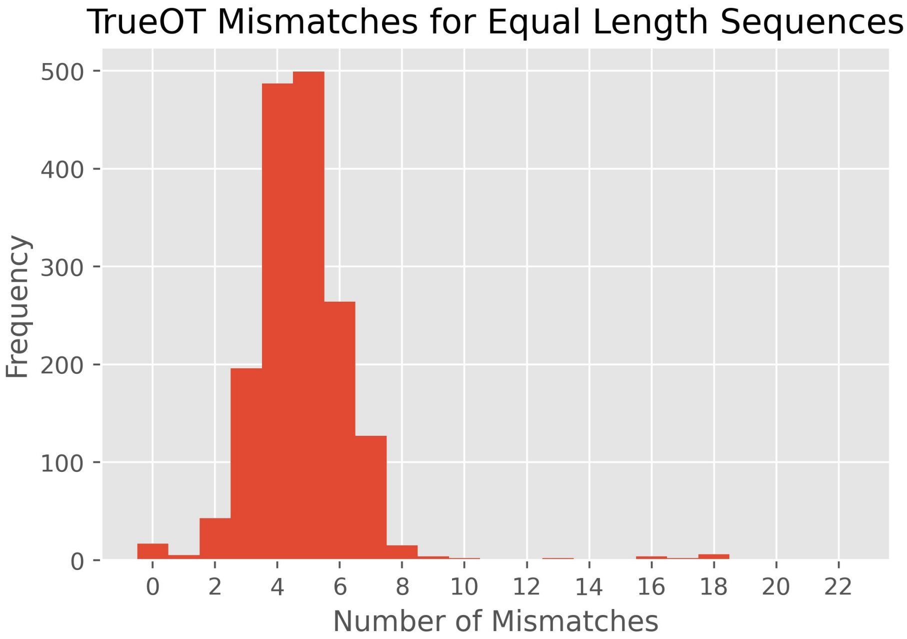


**Figure S1:** The number of mismatches versus bulges depends on the alignment technique used. We used the alignment in the original studies for inclusion in TrueOT and find that the vast majority of sequences of equal length and no “-” provided in the original alignment have a small number of mismatches. A few datapoints exhibit many mismatches, although these may be more appropriately characterized by bulges. We did not re-align sequences in our analysis to avoid the introduction of a new source of variability. In any case, our Siamese models ignore the alignment by removing any dashes and simply left-padding sequences to a fixed length.
